# High-magnitude innovators as keystone individuals in the evolution of culture

**DOI:** 10.1101/242131

**Authors:** Michal Arbilly

## Abstract

Borrowing from the concept of keystone species in ecological food webs, a recent focus in the field of animal behaviour has been keystone individuals: individuals whose impact on population dynamics is disproportionally larger than their frequency in the population. In populations evolving culture, such may be the role of high-magnitude innovators: individuals whose innovations are a major departure from the population’s existing behavioural repertoire. Their effect on cultural evolution is twofold: they produce innovations that constitute a ‘cultural leap’, and, once copied, their innovations may induce further innovations by conspecifics (socially induced innovations), as they explore the new behaviour themselves. I use computer simulations to study the co-evolution of independent innovations, socially induced innovations, and innovation magnitude, and show that while socially induced innovation is assumed here to be less costly than independent innovation, it does not readily evolve. When it evolves, it may in some conditions select against independent innovation and lower its frequency, despite it requiring independent innovation in order to operate; at the same time, however, it leads to much faster cultural evolution. These results confirm the role of high-magnitude innovators as keystones, and suggest a novel explanation for low frequency of independent innovation.

## Introduction

The concept of keystone species, originally suggested by Robert Paine to describe species whose impact on their ecosystem is much greater than their part in it [1–3], has been recently adopted by animal behaviour researchers, to describe individuals whose impact on the population they live in is much greater than their proportion in it, and whose removal from the population would result in a profound and lasting effect on group dynamics [4]. While the general concept is relatively new, effects of such individuals have been noted and documented over decades and across social species, by more situation-specific titles, such as dominants, tutors, or leaders (see detailed review in Modlmeier et al. 2014). Recent studies utilizing the keystone individuals concept have shown, for example, that the presence of a few bold individuals in colonies of social spiders, and the quality of the knowledge these individuals possess affects the colony’s foraging behaviour and success [5,6], and that an ant colony’s nest site selection is faster and more accurate when it includes highly exploratory individuals [7]. The keystone framework is also gaining some traction in conservation biology: it has recently been proposed that identification of keystone individuals and analysis of their effect on the population is valuable in conservation and management of social species [8].

In the context of cultural evolution, we may consider innovators of behaviours that spread in a population, and individuals who serve as a popular copying model, to be keystones [4]. Theoretical and experimental work assessing the role of innovation in cultural evolution has focused on the conditions favouring social learning over innovation (or individual learning) and vice versa (e.g. [9–17]), as well as on the diffusion of innovations [18–20]. Lately, a series of models turned the spotlight onto the way different types of innovations may shape the evolution of culture [21–23]. The different nature that innovations may have pertains to a longstanding dispute in the animal behaviour literature. While it is intuitively clear that not all innovations are similar in their inception and impact, how can we define the differences between them in general terms?

In a recent paper, we approached this issue by describing behavioural innovations as measured by their magnitude [24]. Relying on a previous definition [25], we suggested that any new behaviour, no matter how similar to behaviours already in the populations’ behavioural repertoire, should be considered an innovation; however, we argued that these innovations may differ in how close to or far from the population’s mean behaviour they are. Innovations that are far from the mean are considered high-magnitude innovations, while innovations that are close to the mean are considered of low-magnitude (see detailed discussion in [24]). Offering a great increase in the population’s fitness, high-magnitude innovations that spread in the population may therefore be viewed as a cultural ‘leap’ [21].

High-magnitude innovations, by definition, differ significantly from familiar behaviours. They may include the introduction of a new object to interact with [26], a new territory to forage in [27], a new feeding method to utilize [28], or a new song to replicate [29]. Viewing others interact with an unfamiliar object may allow neophobic individuals to overcome their fear, or simply draw attention to an object that copiers have not noticed before [30,31]. Thus, high-magnitude innovations allow copiers of the innovation to explore a new domain and perhaps modify it by innovating themselves. Models focusing on the effect of different innovation types on human cultural evolution have utilized the latter idea, suggesting to account for the punctuated evolutionary pattern found in the human artifact archeological record [21–23]. High-magnitude innovators may therefore not only serve as keystone individuals by generating cultural leaps, but also by facilitating socially induced innovations, that further modify their own.

In this study, I expand upon our previous work on the magnitude of innovation in social animals [24], to include cultural evolution. I investigate whether a trait allowing socially induced innovation can evolve, examine the effect of such a trait on the evolution of independent innovation and on the magnitude of innovation, and finally, analyze how all these traits interact to shape the progression of culture.

## THE MODEL

I simulated a population of individuals genetically varying in their (1) tendencies to innovate and to copy others; (2) innovation magnitude, and (3) tendency to modify high-magnitude innovations they have copied. A generation’s life began with a series of *T* = 10 discrete learning steps, in each of them individuals acquire one new behaviour either by innovating, or by copying the innovations that others produced during that specific learning step *t* (0 < *t* ≤ T). Individuals who copied high magnitude innovations in step *t* could, based on their genetic tendency, be “inspired” to innovate in the next learning step (*t*+1), to produce a modification of the copied innovation. This modified innovation could be copied by others during that step (*t*+1 ≤ *T*) as any innovation, and could serve as a basis for further socially induced innovations in the next learning step (*t*+2 ≤ T), in the same manner. After the *T* steps of the learning phase, individuals applied the behaviours they have acquired, with greater weight given to higher-paying behaviours. Individuals then produced offspring in proportion to the relative payoff they have accumulated during their lifetime, and died. The mean of the highest paying behaviours learned by parents was defined as their generation’s cultural contribution, and considered the new generation’s behavioural baseline for cultural evolution calculations.

### The population

A population of *n* = 100 individuals is modeled, with each individual characterized by three focal genes: *L* (Learning gene), *I* (Innovation magnitude gene), and *C* (socially induced innovation gene). The learning gene, *L*, determined the probability the individual will, at each learning step, produce an independent innovation, or copy a conspecific’s innovation. There were 11 possible alleles in this gene: 0, 0.1, 0.2 … 1, where 0 coded for full-time copying, or social learning, 1 for full-time independent innovation, and all other alleles for a combination of the two (e.g. a carrier of the 0.3 allele spent 30% of the time, on average, copying, complemented by an average of 70% independent innovation). The innovation magnitude gene, *I*, affected how far from the population’s norm an individual’s innovations will be when innovating. There were again 11 possible alleles in this gene: 0, 0.1, 0.2 …, 1, which represented standard deviations from the population’s mean behaviour; this value was used to draw a value from a normal distribution whose mean was the population’s mean behaviour, and standard deviation was the individual’s *I* allele (see below). The socially induced innovation gene, *C*, determined the probability that, after copying a high-magnitude innovation, the copier will proceed to modify this innovation in its next learning step. This gene included three alleles: *C_0_* – for zero probability, i.e. no effect; *C_sqrt_* – the square root of the individual’s probability to innovate as set by its *L* allele, i.e. an increase in innovation probability that is proportional to genetic tendency for independent innovation; and *C_1_* – for a probability of 1, i.e. the individual is certain to innovate. Just like independent innovation, the magnitude of an individual’s socially induced innovation was determined by its genotype in the *I* gene.

### Learning phase

All individuals in the population had a limited number of learning steps *T* = 10. In each of these steps they acquired one new behaviour, either by innovation or by copying an innovation a conspecific has produced at that specific step (our previous model analyzing the case of *T* = 100 found no significant differences between the two case, see Arbilly and Laland 2017). At the beginning of each step, it was determined for each individual whether it would innovate or copy, based on the probability dictated by its *L* genotype. Individuals who were to innovate generated a new behaviour. The value of this innovative behaviour (i.e. its payoff) was drawn from a normal distribution whose mean was the population’s mean behaviour, and whose standard deviation was the innovator’s allele in the *I* gene; for convenience, the population’s mean behaviour value was set to 0. Then, individuals who were to copy in this learning step, copied the behaviours generated by innovators. All innovations were ranked according to their value, and which innovations would be copied depended on the selectivity of social learning in the population (which was kept constant per population). The selectivity of social learning was controlled using the variable *D*, defined as 1 – [the fraction of demonstrators copied]. When selectivity was high (high *D*) only innovations with the highest value were copied (e.g. when *D* = 0.9 only the top 10% of innovation were copied); as the selectivity of social learning became lower, copying became more random (and was completely random at *D* = 0).

In cases where individuals copied a high-magnitude innovation, defined as an innovation whose value was greater than 1 (putting it at a distance greater than one standard deviation from the population’s mean behaviour), it was determined whether they will modify this innovation in the following learning step, *t*+1, based on their socially induced innovation (*C*) allele. If they were to innovate, the magnitude of their innovation was set by their *I* allele. These individuals produced an innovation at the beginning of step *t*+1 along with independent innovators (described above). However, for these socially induced innovators, the value of their innovation was added to the value of the high-magnitude innovation they copied in their previous learning step (*t*), to yield a new innovation for step *t*+1. This innovation was then ranked along with all innovations and copied by individuals who, in step *t*+1, are copying others, as described above. The choice to have socially induced innovation triggered only by the copying of high-magnitude innovations, rather than the copying of any innovation, was made in order to set these innovations apart from independent innovations (see Discussion).

### Application phase

After acquiring the behaviours, individuals apply these behaviours and will tend to use them with a frequency directly proportional to the payoff they offer. To calculate the proportion of time allotted to each behaviour, and since payoffs can be negative as well as positive, an exponential transformation of the form:

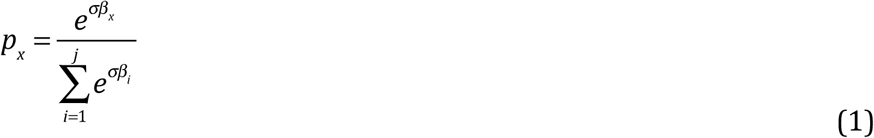
is used, where *p_x_* is the proportion of time spent using behaviour *x, β_x_* is the payoff of behaviour *x*, *i* = 1 … *j* are the behaviours the individual has acquired during its learning phase (*j* = *T*), and *σ* is the application sensitivity: the degree to which agents can distinguish between payoffs in choosing which behaviours to apply. This value is the same for all agents. Following previous analysis [24], *σ* was set to its high value (*σ* = 3.3), such that agents spend a higher proportion of their time applying the highest paying behaviour and little to no time applying low value behaviours. Note that due to the stochastic process used in the simulation to generate new behaviours, then unless there is no innovation in the population, behaviours 1 … *j* will each be unique.

The payoff accumulated from applying the learned behaviours, *W_A_* was then calculated by summing up the multiplications of each behaviour’s payoff and the proportion of time spent applying it:

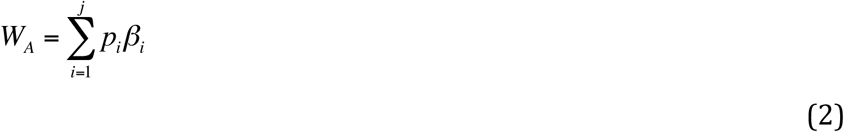

### Selection and reproduction

To calculate the total payoff to individuals in the population, *W_T_*, the payoff obtained both during the learning phase, *W_L_* (which is the sum of all payoffs of behaviours learned), and during the application phase, *W_A_*, was summed using a weight factor *α* = 0.1 to account for the relative time allocated to the learning phase compared to the application phase:

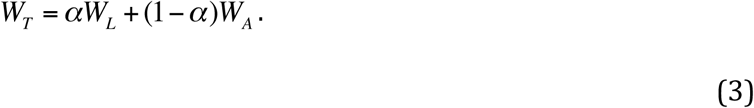

Payoff received for behaviours was included in the learning phase payoff calculation (in the form of *W_L_*) regardless of whether they were applied, as it is assumed that agents perform behaviours when they are learning them, in order to experience their exact payoff.

Individuals then reproduced, producing a number of offspring proportional to their total payoff relative to the payoff of all other individuals in the population. Since the total payoff could be negative, we again use an exponential transformation of the form:

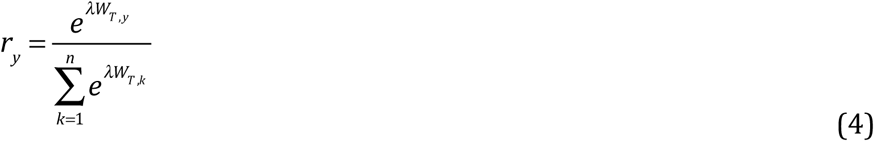

Where *r_y_* is the probability of reproduction for individual *y*, and *λ* is the strength of selection. Following previous analysis [24], *λ* was set to its high value (*λ* = 3.3), to generate strong selection: individuals who obtained higher total payoff had much higher chances to reproduce than individuals who obtained a lower payoff. Among the offspring, mutation occurred at a rate of *μ* = 1/*n* in all genes. Mutation was random and the new variant was drawn from each gene’s pre-defined allele pool.

### Cumulative culture

After parents are selected, each parent’s highest paying behaviour is recorded. The mean of all these behaviours from the parental generation is then counted as that generation’s cultural contribution. This assumption accounts for a situation where a full repertoire of behaviours are transferred to the new generation, and not just one. This mean was then viewed as the new generation’s mean behaviour. Since values of behaviour here are arbitrary, the actual value of this mean does not matter for purposes of innovation in the next generation, and furthermore, using it as the mean for the distribution from which the next generation draws innovations inflates cultural evolution rates, this cultural contribution was set aside and the actual mean used to draw innovations was zero for all generations. These cultural contributions were then used cumulatively to calculate the progress of cultural evolution. For example, if generation 1’s contribution was 1.5, and generation 2’s contribution was 0.5, the final value of culture for generation 2 was 1.5+0.5=2, and so on for following generations. The choice to use the mean of parents’ highest paying behaviours is conservative: using only the single highest paying behaviour for each generation would have resulted in higher cultural rates.

## RESULTS

### Evolution of socially induced innovation

The allele frequency in socially induced innovation gene, *C*, changed with social learning selectivity, *D* (figure 1). The *C_1_* allele, setting the probability of socially induced innovation to 1, had a clear advantage when the selectivity of social learning was low (*D* ≤ 0.1). The allele enhancing the probability of innovation, *C_sqrt_*, was also selected at *D* = 0, although at a much lower frequency, with some advantage over *C_0_* (allele coding for no effect); this advantage of *C_sqrt_* over *C_0_* disappeared when *D* = 0.1. When selectivity was higher, *C_1_* was found at low frequencies, while *C_sqrt_* and *C_0_* appeared at similar frequencies (between 40% and 50% each). It should be noted that in that range of social learning selectivity *D*, independent innovation rate, set by the *L* gene, was close to zero (figure 2a): in most generations individuals had an independent innovation rate of zero, therefore the *C_sqrt_* allele would have no effect on them, similar to *C_0_*.

**Figure 1:**
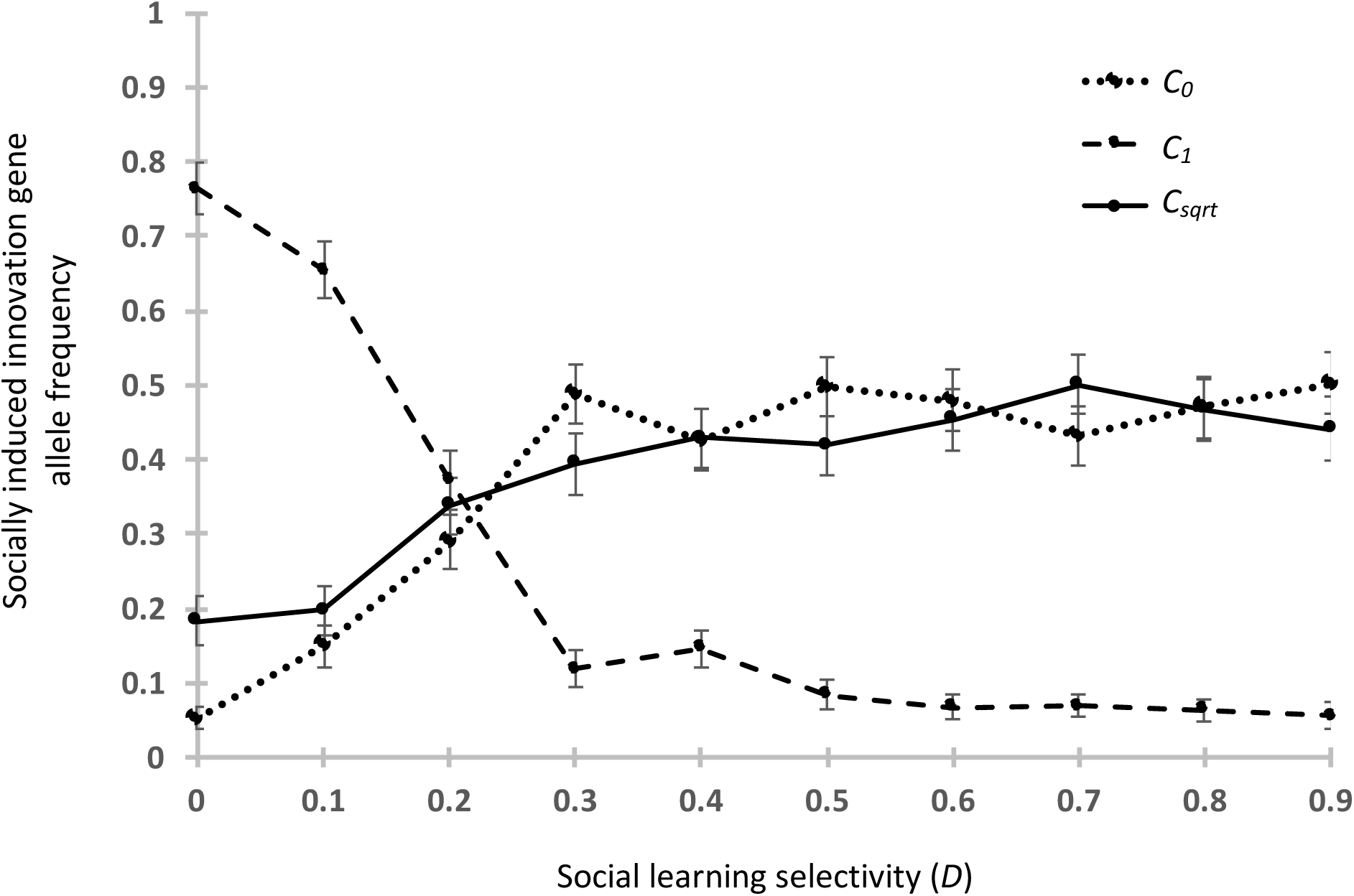
Mean frequency of alleles in the socially induced innovation gene, *C*, as a function of social learning selectivity, *D*. *C_0_* allele codes for no socially induced innovation; *C_1_* allele codes for certain socially induced innovation; *C_sqrt_* sets the probability of socially induced innovation to the square root of the probability of independent innovation (*L* genotype). Means and standard errors are calculated for generations 4001–5000, across 100 repeats of each simulation.

**Figure 2:**
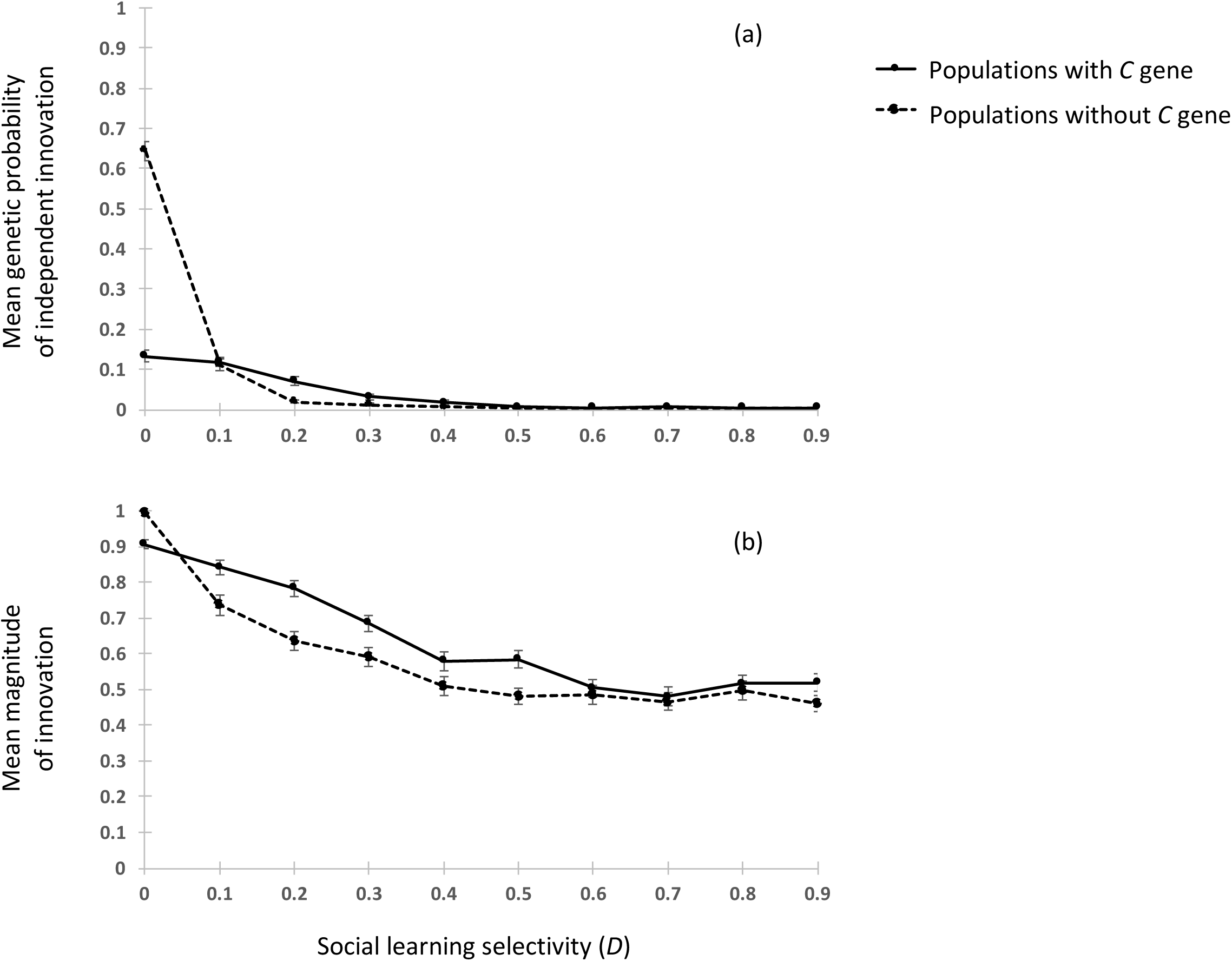
Effect of the socially induced innovation gene, C, on the frequency and magnitude of innovation. (a) Mean frequency of independent innovation, based on mean genotype in the *L* gene; (b) Mean magnitude of innovation, among individuals with the genetic potential of independent and/or socially induced innovation. Means and standard errors are calculated for generations 4001–5000, across 100 repeats of each simulation.

### Rate of independent innovation in the presence of socially induced innovation

A comparison of the genetic probability of independent innovation rate in the presence and in the absence of the *C* gene shows an effect changing with the selectivity of social learning, *D* (figure 2a). While in the absence of C the genetic probability represents the expected probability of innovation in the population, in the presence of C, the actual rate of independent innovation may be lower than the genetic probability, as individuals may use some of their learning steps for socially induced innovation, instead of drawing between innovation and social learning based on their *L* allele. When the selectivity of social learning was at its lowest – where copying is completely random – socially induced innovation significantly decreased the rate of independent innovation. When social learning selectivity was poor while still eliminating the worst innovations (*D* = 0.1), the rate of independent innovation was the same with and without *C*. When selectivity was higher but still in the low range (0.2 ≤ *D* ≤ 0.5), the rate of independent innovation was slightly higher in the presence of the *C* gene. As the effect is very small, and due to the complicated frequency-dependent interaction between the three genes, it is difficult to determine whether this is due to noise created by drift in the *C* gene, because socially induced innovations increase the benefit of independent innovation by increasing the competition, because carriers of the *C_sqrt_* allele benefit when also carrying an *L* allele with a value that is higher than zero, or some combination of these. However, more selective social learning resulted in similar, close to zero rates of independent innovation, with and without socially induced innovation.

### Magnitude of innovation in the presence of socially induced innovation

When the selectivity of social learning was at its lowest – where copying was completely random – the magnitude of innovation was lower in the presence of socially induced innovation, although still very high (0.91 compared to 0.99; figure 2b). In the medium range of social learning selectivity, however, the magnitude of innovation was consistently higher in the presence of socially induced innovation, and as in the absence of socially induced innovation, decreased as selectivity in social learning increased.

### Cultural evolution

Culture as measured by the accumulation of innovations, was higher when the selectivity in social learning (*D*) was lower (figure 3). Socially induced innovation (the *C* gene) increased the rate of cultural evolution; this effect was found even in the case of random copying (*D* = 0), where the rate of independent innovation was much lower in the presence of socially induced innovation than in its absence (figure 2a).

**Figure 3:**
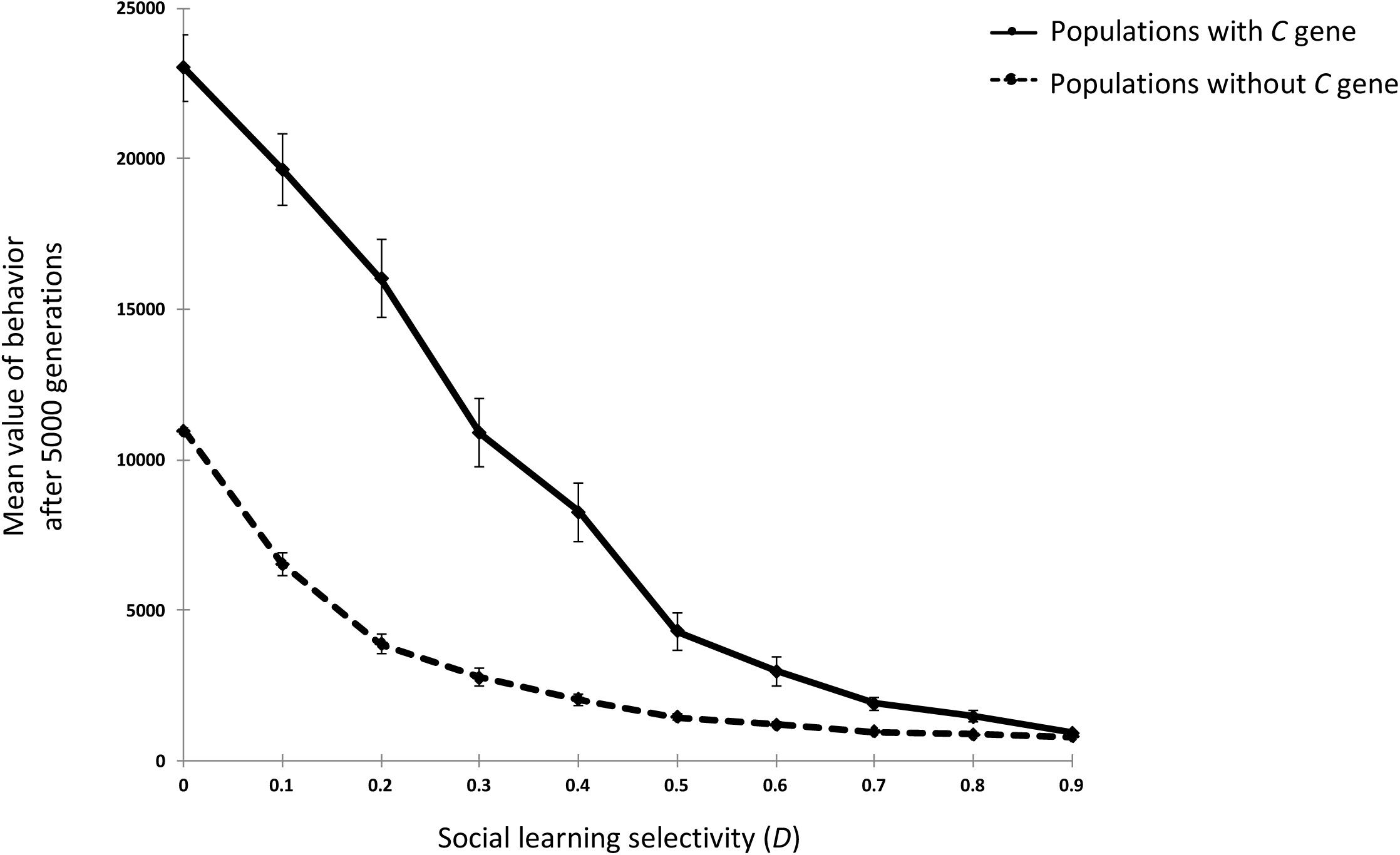
Effect of socially induced innovation gene, *C*, on cumulative culture. Means and standard errors are calculated over 100 repeats of each simulation.

## Discussion

Socially induced innovations would seem to have a clear advantage: building on a known high-magnitude innovation, they offer the possibility of generating an even better innovation, with a lower risk compared to independent innovation. That is, even if the socially induced innovation resulted in a lower value behaviour compared to the independent innovation it was building upon, it is still less likely to be below the population’s mean value of behaviour, unlike independent innovations. Still, socially induced innovations do not evolve when the selectivity of social learning is high: in that situation, others are likely to copy a high-magnitude socially induced innovation, without incurring the possible cost of producing a low-magnitude innovation. The cost for the socially induced innovator here is not only in having a lower value behaviour in its repertoire, but also in missing the chance of copying a better behaviour produced by someone else at that time step. This opportunity cost stems from the assumptions of the model, whereby individuals must perform the behaviour in order to learn it, know its exact payoff, and be “inspired” to modify it further with their own innovation.

Most significant is the effect of socially induced innovation on the rate of independent innovation when copying was random (*D* = 0). In that condition, in the absence of the *C* gene, the rate of independent innovation is up to 0.64±0.02, but when incorporating the *C* gene, the rate was down to 0.13±0.01. The magnitude of innovation was also somewhat lower. The dominating allele in the *C* gene at that time was *C_1_*, guaranteeing a socially induced innovation whenever a high-magnitude innovation was copied. This combination of traits is, perhaps unsurprisingly, “safer” than a high rate of independent innovation alone, for the reason discussed above. It should be noted that, while this result was found when social learning selectivity is low, the selectivity in application of behaviour is high, thus individuals do not blindly utilize behaviours; they are simply unable to judge the value of a behaviour without performing it first themselves. Regardless of the specific condition, it demonstrates how socially induced innovation may affect independent innovation, in a situation where independent innovation would otherwise be highly favoured. While lowering the rate of independent innovation, and the magnitude of all innovations, the *C* gene also lead here to a much higher rate of cultural evolution: socially induced innovations may be copied by others, who may subsequently use them as a basis for further socially-induced innovations, resulting in a cascade of innovations. Altogether, socially induced innovation, which can only act in the presence of high-magnitude independent innovation, selects here against high-magnitude independent innovators, and by lowering their frequency makes their role, as initiators of the innovation cascade, more crucial. In other words, it makes them keystone individuals.

The definition of keystone individuals, as discussed by Modlmeier et al [4], asserts that keystones cannot be “generic”: if removed, their niche cannot simply be filled by others. In the model presented here, individuals may be genetically identical, but few may, by chance, produce a high-magnitude independent innovation, while others may copy it and modify it. Their role as keystones is determined based on the result of their actions. Their independent innovations are a product of probability, and within a generation lifetime do not depend on whether others may have or may not have produced high-magnitude independent innovations of their own. Thus, the removal of a specific keystone individual would indeed not result in another individual in the population producing a high-magnitude innovation in its place.

The results of the model provide, through proof of concept, insight into the co-evolution of independent and socially induced innovation. As human technology is undoubtedly made of cascades of innovations [32], the finding that socially induced innovations may select against independent innovation is highly relevant, and fits nicely with results of models that combine these two types of innovations, to demonstrate how human culture may have evolved in “bursts”, composed of initial ‘lucky leap’ innovations that are followed by further innovations that are inspired by the leap [21–23]. Furthermore, the results presented here demonstrate how socially induced innovation may help maintain independent innovations, or lucky leaps, at a low frequency, when it is tough to gauge the payoff of a behaviour without first-hand experience (see discussion of selectivity above).

While the model aims to be general, cumulative culture in nonhuman animals, to the extent that it exists, is difficult to track. Some exceptions to this rule, however, are bird song [29], and whale song [33], where populations have been documented evolving unique vocal repertoires. Studies in bird song suggest possible costs to song innovation (e.g. the signal not conveying the signaler’s intended information [29]), as well as benefits (adjusting song to new ecological circumstances, e.g. songs that travel better in an urban environment [34–36]). They also suggest that innovations often arise through copying errors [29]. This is especially interesting in the context of socially induced innovation, as a novel song (i.e. an innovation), only performed by a single individual, would seem more likely to be replicated with errors by listeners (i.e. lead to socially induced innovation), compared to a song performed by many in the population (i.e. the mean behaviour).

Is the concept of keystone individuals conducive to our understanding of the evolution of culture? What if, for example, individuals were induced to innovate by copying any innovation, regardless of its magnitude? In such a case, socially induced innovations would have no benefit over independent innovations: if the original innovation they innovate upon is not of high value, socially induced innovations are just as likely to result in below-average behaviour as an independent innovation. Thus, cultural evolution rates with such indiscriminate socially induced innovations is likely to be the same as in their absence. Having the keystone concept in mind contributed much to the design of the model presented in this paper, and in turn, to its insight into the possible evolutionary interaction between independent innovation, socially induced innovation, and innovation magnitude, and how this interaction can shape the evolution of culture.

## Supporting materials

The Matlab code for the simulations used in this paper has been uploaded as part of the electronic supplementary material.

## Acknowldegements

I wish to thank two anonymous reviewers for their thoughtful, constructive comments on a previous version of this manuscript.

